# Determination of the molecular reach of the protein tyrosine phosphatase SHP-1

**DOI:** 10.1101/2020.09.04.283341

**Authors:** Lara Clemens, Mikhail Kutuzov, Kristina Viktoria Bayer, Jesse Goyette, Jun Allard, Omer Dushek

## Abstract

Immune receptor signalling proceeds by the binding (or tethering) of enzymes to their cytoplasmic tails before they catalyse reactions on substrates within reach. This is the case for the enzyme SHP-1 that, upon tethering to the inhibitory receptor PD-1, dephosphorylates membrane substrates to suppress T cell activation. Precisely how tethering regulates SHP-1 activity is incompletely understood. Here, we use surface plasmon resonance to measure binding, catalysis, and molecular reach for PD-1 tethered SHP-1 reactions. We find that the reach of PD-1—SHP-1 complexes is dominated by the 13.0 nm reach of SHP-1 itself. This is longer than an estimate from the structure of the allosterically active conformation (5.3 nm), suggesting that SHP-1 explores multiple active conformations. Using modelling, we show that when uniformly distributed, PD-1—SHP-1 complexes can only reach 15% of substrates but this increases to 90% when they are co-clustered. When within reach, we show that membrane recruitment increases the activity of SHP-1 by a 1000-fold increase in local concentration. The work highlights how molecular reach regulates the activity of membrane-recruited SHP-1 with insights applicable to other membrane-tethered reactions.

**Significance statement:** Immune receptors transduce signals by recruiting (or tethering) cytoplasmic enzymes to their tails at the membrane. When tethered, these enzymes catalyse reactions on other substrates to propagate signalling. Precisely how membrane tethering regulates enzyme activity is incompletely understood. Unlike other tethered reactions, where the enzyme tethers to the substrate, the substrate in this case is a different receptor tail. Therefore, the ability of the receptor-tethered enzyme to reach a substrate can be critical in controlling reaction rates. In this work, we determine the molecular reach for the enzyme SHP-1 and the receptor PD-1 to which it can tether, and show how molecular reach controls receptor signalling.

## Introduction

Immune receptor signal transduction proceeds by the recruitment of cytoplasmic enzymes to their unstructured cytoplasmic tails before they catalyse reactions on other membrane substrates (1–3). A well-studied example is the inhibitory checkpoint Programmed cell death protein 1 (PD-1) on T cells that contains immunoreceptor tyrosine-based inhibition (ITIM) and switch (ITSM) motifs (4). Ligand binding induces phosphorylation of these motifs that can then recruit the tyrosine phosphatase SHP-1 by its SH2 domains. When tethered to PD-1, SHP-1 can inhibit T cell activation by dephosphorylating the T cell receptor and the co-stimulatory receptor CD28 (5–8) (Fig. 1A). Precisely how the activity of SHP-1 is regulated by membrane recruitment is incompletely understood.

**Figure 1:**
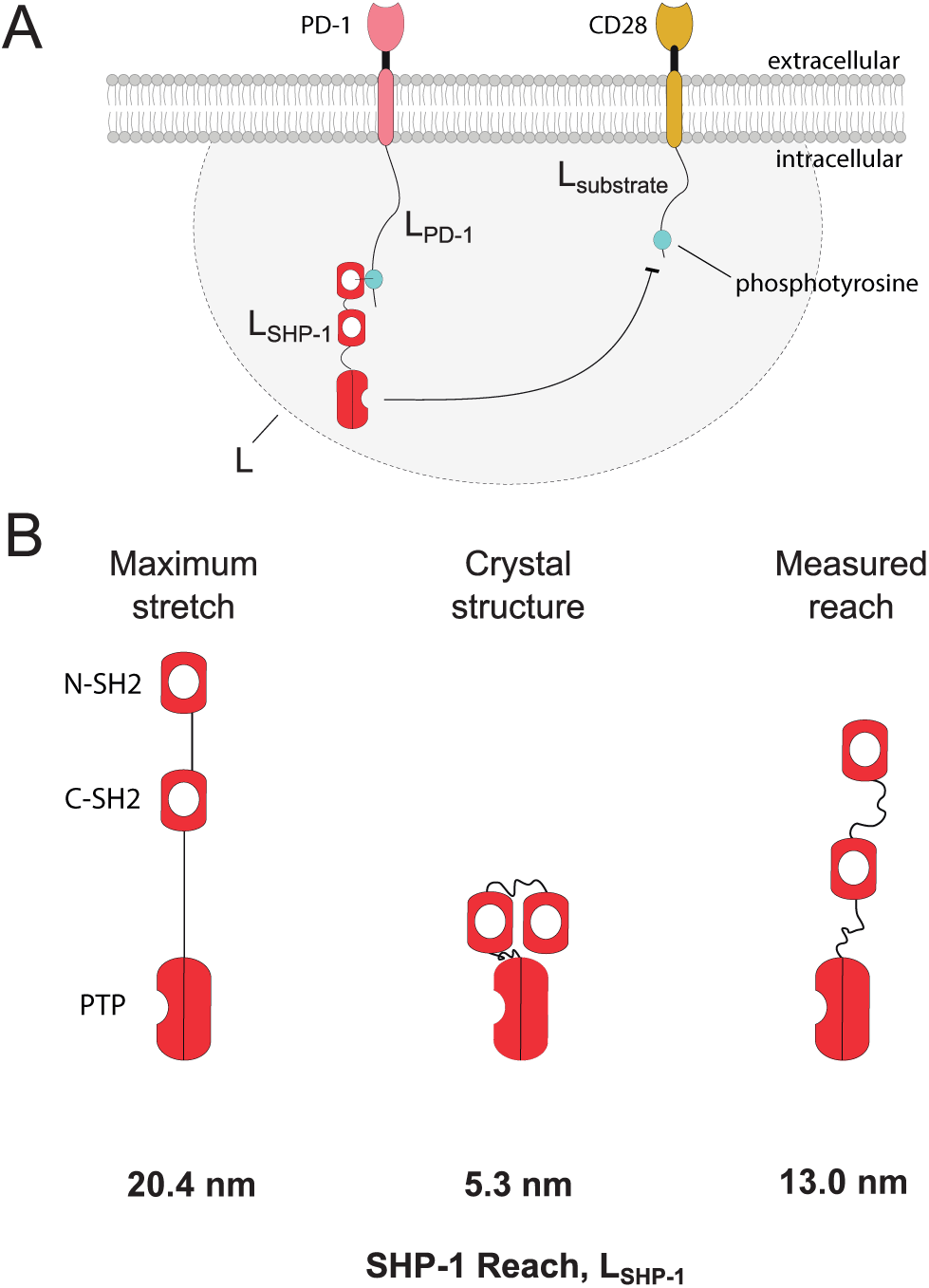
Molecular reach in immune receptor signal transduction. (A) Schematic of a tethered de-phosphorylation reaction mediated by the tyrosine phosphatase SHP-1 (red) recruited to the membrane by the inhibitory receptor PD-1 (pink) acting to dephosphorylate the co-stimulatory receptor CD28 (orange). The molecular reach of the reaction, *L* (gray area), is determined by the molecular reach of PD-1 (*L*_PD−1_), CD28 (*L*_substrate_), and SHP-1 (*L*_SHP−1_). It determines whether the substrate is within reach (within gray area) and the local concentration of SHP-1 when this is the case. (B) Estimates of SHP-1 molecular reach (*L*_SHP−1_) based on sequence (maximum stretch), crystal structure, and experimental measurement in the present work.

Biochemical and structural studies have clearly demonstrated that engagement of the SH2 domains of SHP-1, and its family member SHP-2, induces a conformational change from a closed low-activity state into an open high-activity state (9–18). When quantified, this binding-induced allosteric activation can increase catalytic rates by up to 80-fold (9) and therefore, this is a mechanism by which membrane recruitment can regulate enzyme activity and has motivated the development of therapeutic allosteric inhibitors (15).

In addition to allosteric activation, membrane recruitment also serves to tether SHP-1 in a small volume increasing the local concentration of SHP-1 experienced by substrates (3). Tethering is prevalent in cellular signalling (19) and experimental and mathematical work has shown that it can dramatically increase local concentrations, and hence reaction rates (16, 20, 21), and it can also override enzyme specificity (22). How tethering impacts the local concentration of SHP-1 is presently unknown.

In contrast to previously studied tethered reactions (20, 21, 23), where the enzyme tethers directly to the substrate, the situation is more complicated for immune receptors because the enzyme tethers to a receptor but acts on a different membrane substrate, which is often a different receptor tail (Fig. 1A). Therefore, the potentially high local concentration that results from membrane recruitment may only be experienced by the small subset of substrates within reach. The molecular reach of the reaction (*L* in Fig. 1A) (24) is a biophysical parameter that determines both the fraction of substrate that is within reach and when this is the case, the local concentration (approximately *σ*\* = 1*/L*^3^). Therefore, quantifying molecular reach is critical for understanding the impact of SHP-1 membrane recruitment.

The molecular reach of these reactions is presently unknown. There are three molecules that contribute to the reach: the receptor tail, the enzyme, and the substrate (24). Polymer models, such as the worm-like chain (WLC), have been successfully in estimating reach for unstructured polypeptide chains based on the contour (*l*_*c*_) and persistence (*l*_*p*_) lengths as *L*_peptide_ = (*l*_*c*_*l*_*p*_)^1*/*2^ (20). This theoretical approach predicts a reach of *L*_PD-1_ = 3.0 nm for the ITSM of PD-1 located 55 amino acids from the membrane (using *l*_*c*_ = 55 × 0.4 nm, where 0.4 nm is the contribution of each amino acid and *l*_*p*_ = 0.4 nm for random amino acid sequences (20, 25)). In the absence of other contributions, this reach is comparable to the dimensions of receptor extracellular domains implying that surface receptors must come into (or nearly into) contact to enable reactions.

However, SHP-1 may significantly contribute to this reach. SHP-1 tethers with its dominant N-terminal SH2 domain and catalyses reactions with a C-terminus protein tyrosine phosphatase (PTP) domain (9, 11, 16) as shown in Fig. 1A. Based on the crystal structure of the allosteric open conformation of SHP-1 (14), the reach between the N-SH2 and the catalytic pocket is estimated to be 5.3 nm. However, SHP-1 may dynamically explore conformations not observed in crystals to achieve a longer reach. A potential upper bound can be estimated by assuming that all linkers are maximally stretched obtaining a reach of 20.4 nm (Fig. 1B). However, the combination of structured domains, flexible linkers, and specific interactions between them makes it difficult to accurately predict the reach of multi-domain proteins like SHP-1.

Here, we extend a previously described surface plasmon resonance (SPR) based assay (16) to measure binding, catalysis, and molecular reach for PD-1 tethred SHP-1 reactions at 37°C. We find a reach of 6.55 nm for PD-1 and a reach of 13.0 nm for SHP-1, suggesting it dynamically explores a range of open conformations. The molecular reach shows that membrane-recruitment can increase the activity of SHP-1 by a 1000-fold increase in local concentration, which is larger than the activity increase by allostery, and that clustering is required for PD-1–SHP-1 complexes to reach the majority of substrates. The work highlights the role of molecular reach in regulating the activity of tethered SHP-1 reactions providing insights widely applicable to immune receptors.

## Results

### Extended SPR assay determines biophysical parameters for tethered reactions by SHP-1

We previously described an SPR-based assay and fitting procedure for extracting biophysical parameters of tethered reactions (16) (Fig. 2A). In that assay, murine SHP-1 was injected over initially phosphorylated ITIM peptide from murine LAIR-1 coupled to polyethylene glycol (PEG) repeats at 10°C and binding of SHP-1 (via its SH2 domains) is monitored by SPR over time. Since SH2 domains bind phosphorylated peptides, it was observed that although binding initially increases, it rapidly decreased over the injection time because of peptide tether dephosphorylation primarily by tethered reactions (when SHP-1 is bound to the tether at the surface and acting on another tether) with some contribution from solution reactions (when SHP-1 directly acts on a surface tether from solution without binding to a tether). The resulting multiphasic SPR traces were fit by a partial-differential-equation mathematical model describing multi-particle densities (MPDPDE model) including densities, spatial auto-correlation, and pair-correlations between tethers.

**Figure 2:**
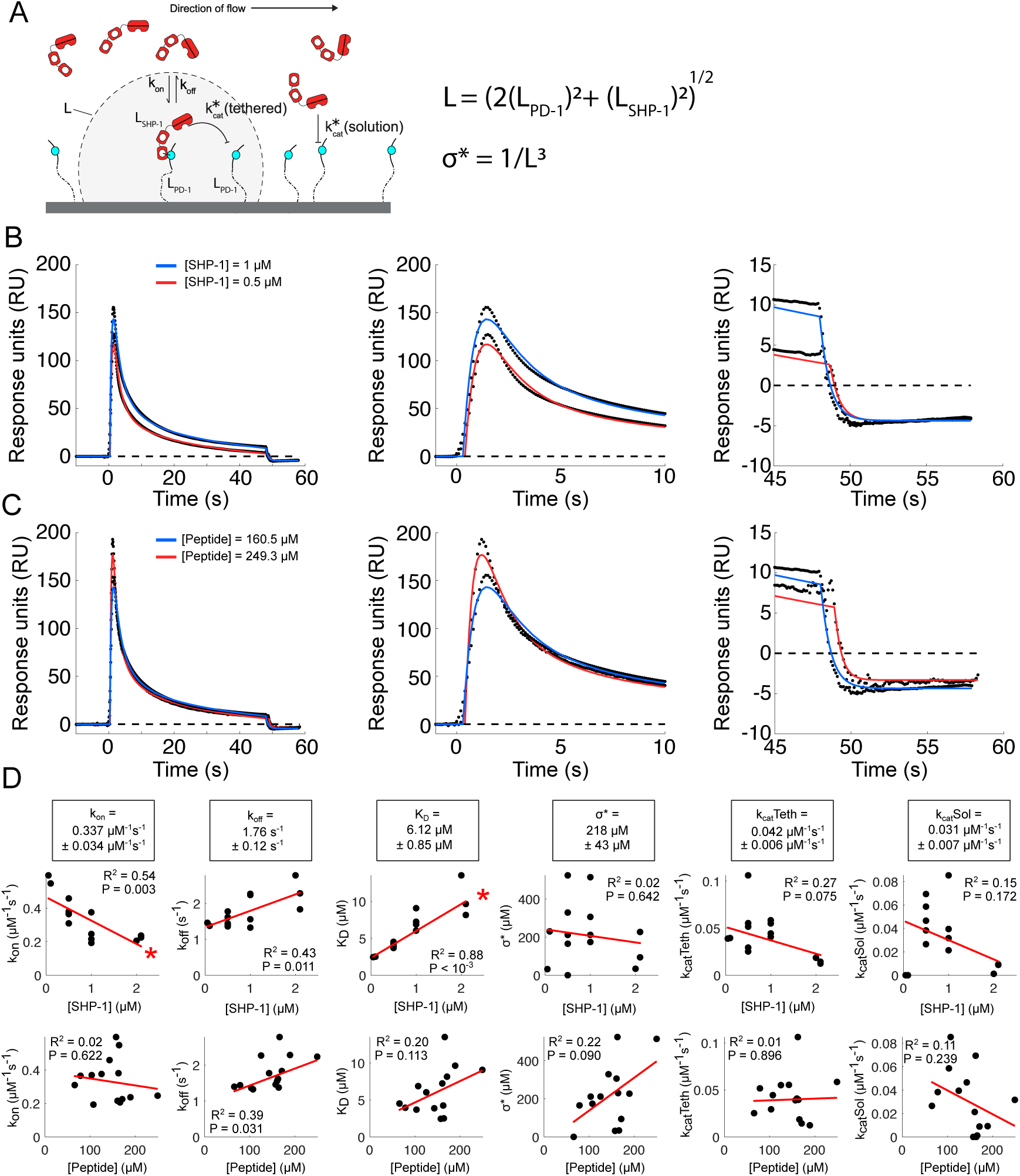
Extended SPR-based assay for tethered catalytic reactions can recover biophysical parameters independent of experimental conditions at 37°C. (A) Schematic of the SPR assay where SHP-1 (analyte) is injected over immobilised phosphorylated PEG28-PD-1 peptides. (B,C) Representative SPR traces (black dots) and MPDPDE model fit (solid lines) for two representative (B) injected human SHP-1 concentrations and (C) two immobilised PEG28-PD-1 concentrations. Middle and right panels show early and late time data, respectively. (D) Fitted parameters (black dots) against SHP-1 concentration (top row) and PEG28-PD-1 concentrations (bottom row) with linear regression (red line; *R*^2^ and p-values without corrections). Red asterisks denote significant correlations at 5% level for Bonferroni-corrected p-values. Averages and SEMs of fitted parameters are shown in boxes above corresponding plots (n=14). All parameters are summarised in Table S1.

Here, we focus on human SHP-1 interacting with a phosphorylated ITSM peptide from human PD-1 initially coupled to 28 repeats of PEG (PEG28-PD1) and data is collected for different injected SHP-1 and immobilised PEG28-PD-1 concentrations at 37°C (Fig. 2). In contrast to our previous experiments (16), we routinely noted a difference in the baseline SPR signal between the start at 0 seconds and the end at 45 seconds of the SHP-1 injection, for example visible in Fig. 2B,C. Using alkaline phosphatase, we found that this difference is not a result of phosphate mass being lost from the surface by dephosphorylation (Fig. S1) but can be explained by nonspecific binding of the enzyme (Fig. S2). We therefore extended our previous MPDPDE model to capture nonspecific binding by including three additional parameters (*p*_start_, *p*_stop_, *p*_*nsb*_), see Methods for details.

We find that this extended eight-parameter MPDPDE model (*k*_on_, *k*_off_, *k*_cat_(tethered), *σ*\*, *k*_cat_(solution), *p*_start_, *p*_stop_, *p*_*nsb*_) closely fits the SPR data (Fig. 2A-C). We perform Markov Chain Monte Carlo analysis to assess if the parameters can be uniquely identified and find this to be the case (Fig. S3). For each SPR trace, we use this extended MPDPDE model to extract all eight parameters, including the molecular reach (or, equivalently, the local concentration *σ*\* = 1*/L*^3^).

### Biophysical parameters associated with catalysis are independent of experimental conditions

We next determined whether the biophysical parameters can be identified independently of experimental variables, specifically the SHP-1 and PEG28-PD-1 concentrations (Fig. 2D). We plot each fitted biophysical parameter as a function of the experimental variables finding that parameters associated with catalysis (*k*_cat_(tethered), *k*_cat_(solution), *σ*\*) are independent of these variables (correlations are not significant). In contrast, the binding parameters (*k*_on_, *k*_off_, and *K*_D_ = *k*_off_*/k*_on_) exhibited a correlation with SHP-1 concentration, with a statistically significant correlation for *k*_on_ and *K*_D_ after a correction for multiple hypotheses is implemented (indicated by red asterisks in Fig. 2D). A possible explanation for this correlation could be that higher concentrations of SHP-1 lead to steric crowding effects on the surface, whereby volume exclusion reduces the ability for more SHP-1 molecules to bind to the surface reducing apparent binding. Based on these results, we conclude that the SPR assay and subsequent model fitting procedure can be used to determine catalytic rates and molecular reach independent of the SHP-1 and the immobilised peptide concentrations.

### Isolating the molecular reach of SHP-1 by varying the tether length

The molecular reach of the reaction, *L* = (*σ*\*)^−1*/*3^, determined using the SPR-based assay involves two components: the reach of the PEG-peptide tether and the reach of the enzyme. As the reach contributed by the tether is progressively decreased (e.g. by using progressively shorter tethers), eventually the molecular reach of the reaction will be wholly determined by the reach of the enzyme. Indeed, if we assume that the reach of the tethers and enzyme can be effectively modelled by worm-like chains, an equation can be derived to relate *L* with the contour length of the tether (Eq. 4, see Methods). This model predicts that the squared molecular reach of the reaction should be linearly related to the length of the tether (Eq. 5), with the reach of the enzyme being the vertical intercept (i.e. when the tether length is nil).

Therefore, we performed the SPR-based assay using a different number of PEG repeats (*N*_*P EG*_ = 0, 3, 6, 12, 28) coupled to the same short PD-1 ITSM peptide (Fig. 3A). As before, the extended MPDPDE model was able to fit the data and produced binding and catalysis parameters that were similar for different length PEG linkers with the exception of *σ*\*, which progressively increased as the number of PEG linkers were reduced (Fig. 3B). This is expected because with shorter PEG linkers the local volume that SHP-1 is confined to decreases, thereby increasing local concentration.

**Figure 3:**
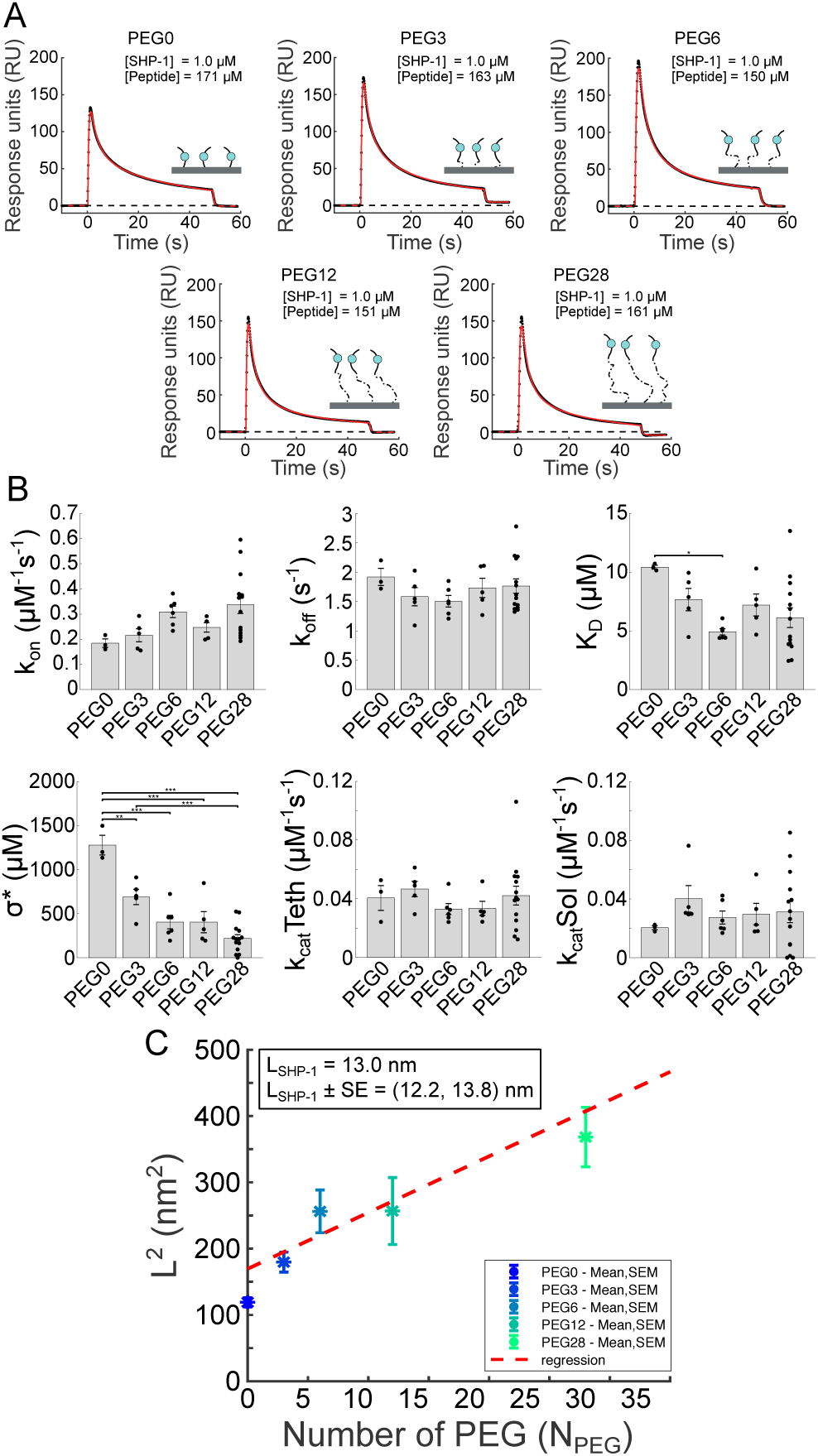
Isolating the molecular reach of SHP-1 by varying PEG-PD-1 tether lengths. (A) Representative SPR traces (black dots) and extended MPDPDE model fits (red lines) for the indicated number of PEG linkers (*N*_PEG_ =0, 3, 6, 12, 28). (B) Averages and SEMs for fitted parameters at indicated PEG linker length. Individual data points are plotted as black dots. Pairwise multiple t-test of parameters and PEG lengths shown for (*) 0.05, (**) 0.01, and (***) 0.001 significance. Significant differences are largely observed for *σ*\*, which determines the molecular reach of the reaction (*L* = (*σ*\*)^−1*/*3^). All parameters are summarised in Table S1. (C) Average squared molecular reach of reaction plotted against number of PEG linkers. Red dashed line indicates regression (p-value = 10^−11^; see Methods). The indicated molecular reach of SHP-1 is estimated by the vertical intercept using the regression line.

As expected, the squared molecular reach of the reaction (determined by converting *σ*\* to L) increased with the number of PEG linkers (Fig. 3C). Using regression, we determined the vertical intercept, and hence the molecular reach of SHP-1, to be *L*_SHP-1_ = 13.0 ± 0.8 nm. This value is between estimates obtained using crystal structure and maximum stretch (Fig. 1B).

We also performed experiments in which the PD-1 peptide is directly coupled to the surface without any PEG linkers. In Fig. 3C, this is labeled PEG0 and produced a molecular reach of 10.9 ± 0.3 nm. Although this value is also within theoretical estimates and similar to the value obtained by the intercept method above, we reasoned that it may be less accurate because this very short peptide can introduce steric hindrance to binding and catalysis, which is reflected in the larger value of *K*_D_ and smaller value of *k*_cat_(solution) that this peptide produces compared to peptides with PEG linkers, and this effect was also noted in our previous work (16).

### PD-1 contributes less than SHP-1 to the molecular reach of the reaction

Given that the molecular reach of the reaction is determined by both the enzyme and tether, we next sought to determine the molecular reach of the cytoplasmic tail of the surface receptor. We injected SHP-1 over immobilised peptide corresponding to the cytoplasmic tail of PD-1 from the membrane to the ITSM. This N-terminally biotinylated peptide contained 64 aa with the phosphorylated tyrosine in the ITSM being 55 aa from the membrane (position 248 in the native sequence). The extended MPDPDE model was fit to the SPR traces (Fig. 4A) and provided estimates of the biophysical parameters (Fig. 4B).

**Figure 4:**
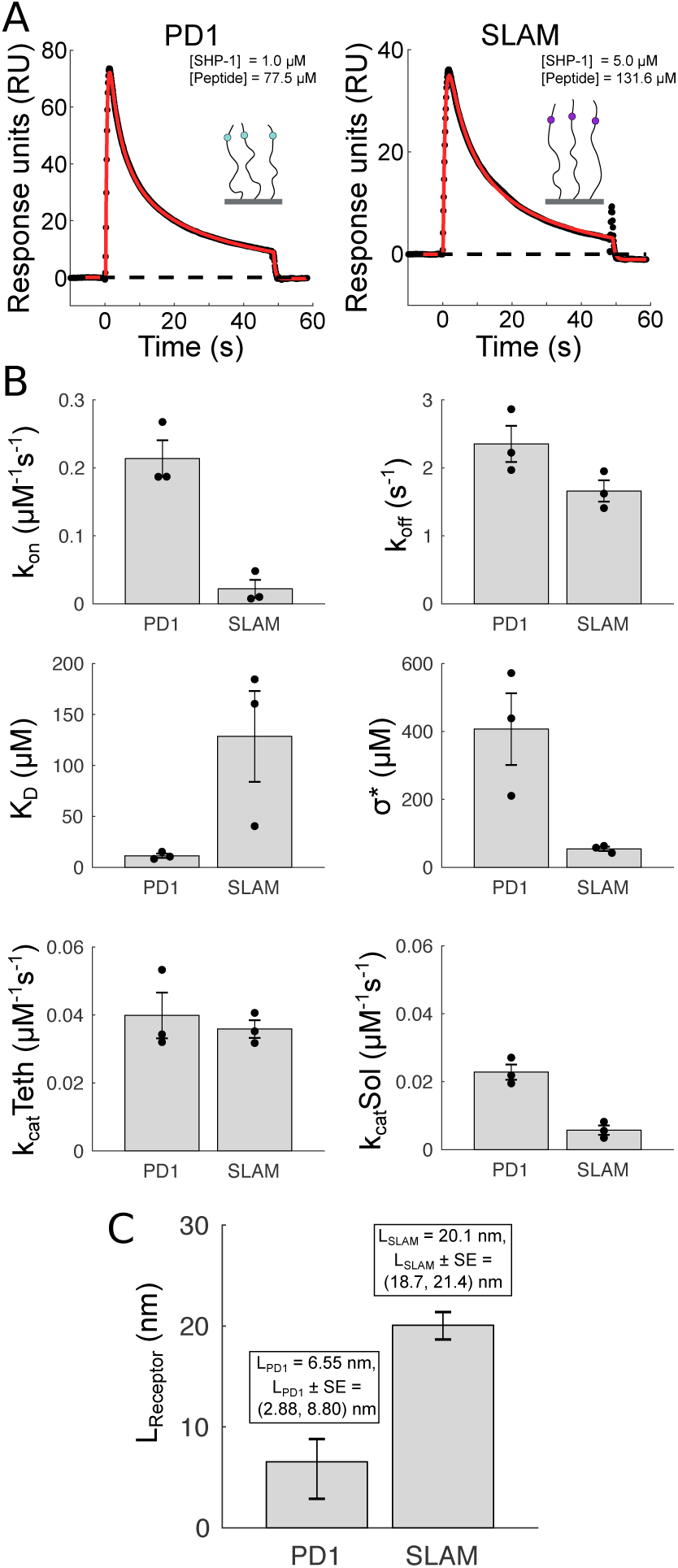
Contribution of PD-1 and SLAM cytoplasmic tails to the molecular reach of the reaction. (A) Representative SPR traces (black dots) and extended MPDPDE model fits (red lines) for the singly phosphorylated PD-1 (55 aa to phosphorylated tyrosine) and SLAM (69 aa to phosphorylated tyrosine) peptides. (B) Averages and SEMs for fitted parameters. Individual data points plotted as black dots. PD-1 exhibits a larger local concentration (*σ*\*) consistent with a shorter molecular reach. All parameters are summarised in Table S1. (C) Average molecular reach (±SE) for PD-1 and SLAM calculated by parsing out the reach of SHP-1 using 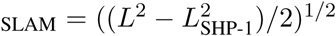, where *L* is the molecular reach of the reaction calculated from *σ*\* in panel B and *L*_SHP-1_ = 13.0 nm.

Using the value of *σ*\*, we calculated the combined molecular reach of the reaction for PD-1-bound SHP-1 acting on PD-1 to be 16 nm. Given that we already obtained an estimate for the reach of SHP-1, we were able to back calculate the reach of PD-1 (see Eq. 4) to be 6.55 nm (Fig. 4C). Thus, we find that PD-1 contributes less to the overall molecular reach of the reactions compared to the SHP-1 reach contribution of 13.0 nm.

We note that the worm-like-chain model would predict a 3.0 nm reach for the PD-1 peptide we have used assuming a persistence length of 0.4 nm that applies to random amino acid chains (20, 25). Therefore, the experimentally measured reach of PD-1 appears to be twice that predicted by the worm-like-chain model, suggesting a preference for extended conformations of this peptide.

The binding affinity between SHP-1 and singly phosphorylated PD-1 was determined to be 11±2 *µ*M.

Using a different assay, Hui et al (5) reported an affinity of 4.28 *µ*M. The ∼2-fold higher affinity they report is likely a result of using a doubly phosphorylated PD-1 peptide.

The ratio of *k*_cat_(tethered) to *k*_cat_(solution) provides an estimate for the strength of allosteric activation of SHP-1 upon SH2-domain binding to PD-1. We find a modest 2-fold increase in activity from this effect (Fig. 4B). Since larger fold increases have been reported previously (9, 11, 16), we further explored this finding. First, we used a standard solution assay whereby SHP-1 acted on a low molecular weight synthetic substrate and confirmed that catalytic activity increased only 2-fold upon addition of a phosphorylated PD-1 peptide (Fig. S4). Second, we previously reported a larger allosteric activation for murine SHP-1 binding to the inhibitory receptor LAIR-1 but at 10°C and therefore performed experiments at this lower temperature, finding again only a modest increase in activity (Fig. S5). We conclude that human SHP-1 exhibits only modest allosteric activation upon binding to singly phosphorylated PD-1.

As a positive control to ensure our SPR-based assay is sensitive to reach, we repeated the experiments using the longer cytoplasmic tail of signalling lymphocyte activation molecule (SLAM), a surface receptor that is also known to recruit SHP-1 (26). The N-terminally biotinylated peptide contained 77 aa with the phosphorylated tyrosine in the ITSM being 69 aa from the membrane (position 327 in the native sequence). Performing the analysis as for PD-1, we find that the molecular reach contributed by SLAM is 20 nm (Fig. 4). This is markedly more than the reach of SHP-1 and comprises 72% of the predicted contour length for the SLAM peptide (*l*_*c*_ ∼ 69 0.4 nm = 27.6 nm). This suggests that SLAM has a larger persistence length than would be expected for random amino acids and/or is otherwise biased towards extended conformations.

Interestingly, we observed a larger 6.2-fold allosteric activation for SHP-1 interacting with SLAM (Fig. 4B) and this is highlighted when plotting the ratio of *k*_cat_(tethered) to *k*_cat_(solution) across all experimental conditions (Fig. S6). However, this allosteric activation for SLAM was a result of a lower *k*_cat_(solution), not a higher *k*_cat_(tethered), compared to PD-1. We also observe a much smaller on-rate for SHP-1 binding to SLAM compared to PD-1. A possible explanation for both observations is that the SLAM peptide may have fewer configurations in which the phosphotyrosine is available for interaction with SHP-1 when in solution.

### Control of surface receptor signalling by the molecular reach of SHP-1

We next used a mathematical model to explore how molecular reach regulates the activity of SHP-1 upon membrane recruitment to PD-1 (Fig. 5A). Using their individual measured reach, we calculated the combined reach of PD-1–SHP-1 complexes as 14.6 nm 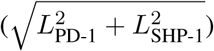. Using this number, we first consider the effective concentration of SHP-1 that a substrate would experience when PD-1–SHP-1 complexes are randomly distributed on the membrane. We find that at physiological densities of PD-1, this effective concentration is ∼1000 *µ*M (Fig. 5B,C), which is ∼1000-fold larger than the ∼1 *µ*M concentration of SHP-1 in the cytosol assuming it is uniformly distributed (16, 27).

**Figure 5:**
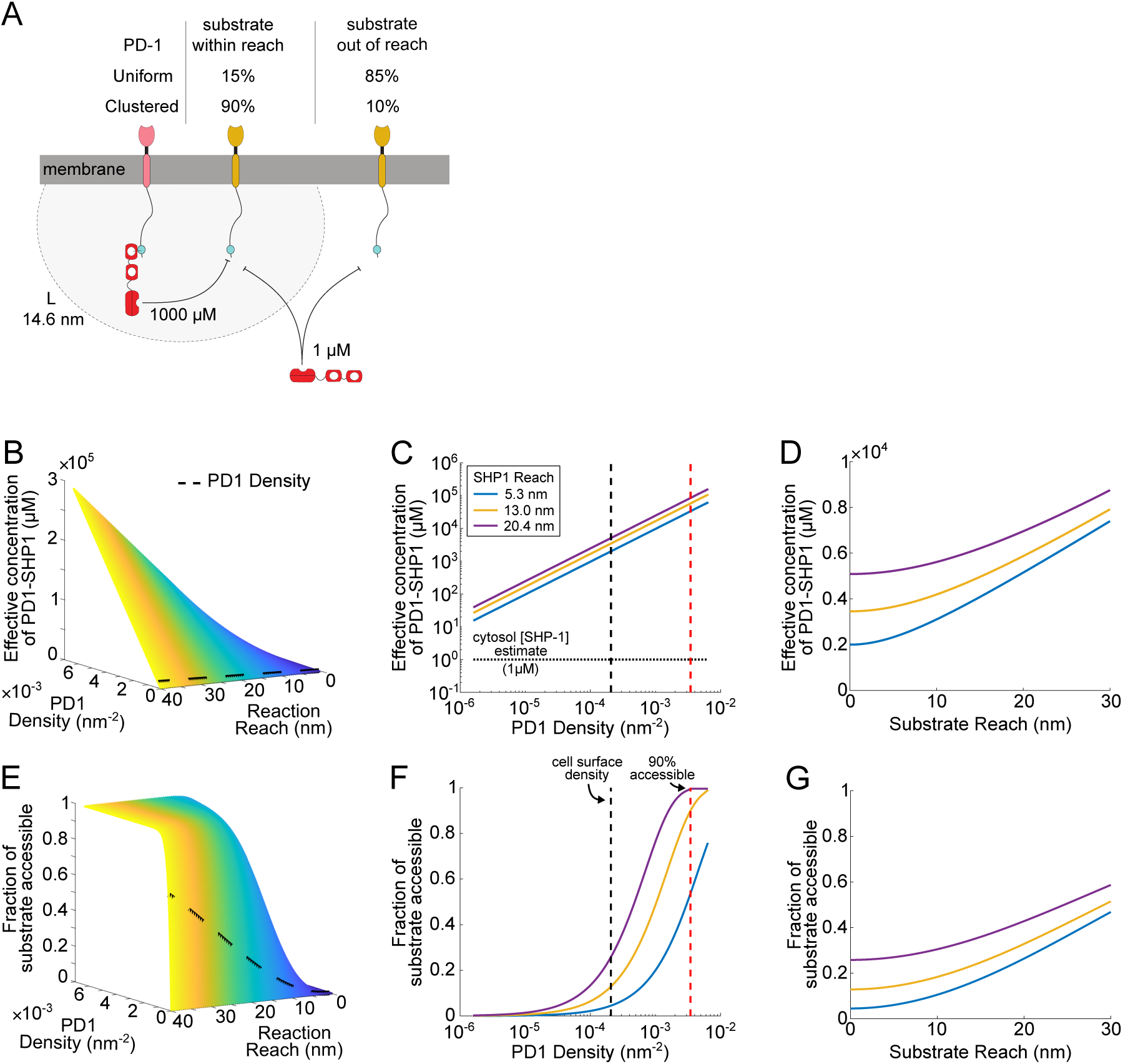
Tethering of SHP-1 to PD-1 generates high membrane concentrations but poor coverage unless PD-1 co-clusters with substrates. (A) Schematic of SHP-1 reactions with substrates, demonstrating tethered reaction (left SHP-1 molecule) and solution reaction (right SHP-1 molecule). (B-D) Effective concentration of PD-1–SHP-1 complex experienced by a substrate and (E-G) fraction of substrate within reach by PD-1–SHP-1 complexes under different conditions: (B,E) Versus the molecular reach of reaction and the density of PD-1–SHP-1 complex on the membrane. The cell surface PD-1–SHP-1 density estimate is shown with black dashed line. (C,F) Versus PD-1–SHP-1 density for different estimates of SHP-1 molecular reach, and fixed PD-1 and substrate molecular reach (6.55 nm and 0 nm, respectively). The density of PD-1– SHP-1 complexes based on a uniform distribution is shown with a black dashed line in B,C,E, and F. Density required to reach 90% of substrate (0.0034 nm^−2^) is shown with red dotted line in F. Estimate of cytosol SHP-1 concentration (1 *µ*M) is shown with dotted horizontal line in C. (D,G) Versus substrate reach (for fixed reach of PD-1 and indicated reach of SHP-1) and PD-1–SHP-1 uniform density (∼2· 10^−4^ nm^−2^). Coloured lines in C,D,F,G refer to the theoretical and experimental estimates of the molecular reach of SHP-1 (Fig. 1B), see legend in C.

In these tethered reactions, even though the effective concentration can be large, the coverage can in principle be low because a random or uniform distribution of surface receptors can allow some substrates to be out of reach (Fig. 5A). We therefore calculated the fraction of substrates that can be accessed by PD-1– SHP-1 complexes for different values of the molecular reach of the reaction and PD-1 density (Fig. 5E,F). If PD-1–SHP-1 complexes are uniformly distributed on the cell surface, we estimate that they are only able to achieve a low ∼15% coverage of the substrates (Fig. 5E,F). Clustering of PD-1 with its substrates, which has been experimentally observed (5, 28), would lead to a higher local density and could therefore be important to improve both the effective concentration and coverage. We find that a clustered density of 0.0034 nm^−2^ (about 10-fold higher than a uniform estimate) is required to achieve a 90% coverage.

We next explored the contribution of the substrate reach to both concentration and coverage. Previously, we noted that the number of amino acids between the membrane and activating or inhibiting tyrosine motifs differed with a median of 33 aa (or 13 nm) or 65 aa (or 26 nm), respectively (16). We therefore repeated the calculations by increasing the contribution of the substrate reach from 0 nm (used in Fig. 5B,C,E,F) to a maximum of 30 nm and found a gradual increase in effective concentration (Fig. 5D) and coverage (Fig. 5G). Within this realistic range of substrate reaches and with uniform distributions, it was not possible to achieve high coverage (e.g. 90%), suggesting that PD-1 co-clustering with substrate is critical for effective inhibition.

## Discussion

Using an extended SPR-based assay for tethered signalling reactions (16), we provide the first estimates of the molecular reach for PD-1 tethered SHP-1 reactions. These values have several implications for how membrane recruitment regulates the activity of SHP-1.

### Impact of membrane recruitment

The membrane activity of SHP-1 is thought to be regulated by allosteric activation. This has been demonstrated using solution catalytic assays whereby engagement of the SH2 domain can increase its catalytic rate (9, 11, 16). However, these solution assays omit tethering, which in the case of immune receptors, determines not only the local concentration but also whether the enzyme can reach its substrates (3, 24). We found that tethering increases the concentration of SHP-1 from ∼1 *µ*M in solution (cytosol) to over ∼1000 *µ*M when tethered (membrane) but importantly, clustering is necessary for the majority of substrates to experience this high local concentration. Interestingly, this 1000-fold increase is much larger than the 2-fold increase in catalytic rate we have measured by allosteric activation when the SH2 domain of SHP-1 is engaged. Therefore, the increase in local concentration can be as important, or possibly more important, than allosteric activation in regulating the activity of SHP-1 at the membrane.

### Implications for allosteric model

The two-state allosteric activation model of SHP-1 (29, 30) is based on crystal structures showing a closed autoinhibitory conformation, where the N-SH2 domain blocks the catalytic pocket (13), and an open conformation, where the N-SH2 is rotated, exposing the catalytic pocket (31). Interestingly, the molecular reach of SHP-1 that we report when tethered in the higher activity state (13.0 nm) is longer than the reach obtained from the open conformation structure (5.3 nm). This suggests that SHP-1 utilises flexible linkers to achieve a spectrum of open states with a longer reach.

### Relation to SHP-2

Although previous reports have demonstrated that allosteric activation of SHP-1 can be induced by singly phosphorylated peptides engaging a single SH2 domain (9, 11, 16), recent reports have suggested that allosteric activation of SHP-2 requires simultaneous binding of both SH2 domains on the same (18) or across different PD-1 peptides (17). Although SHP-1 in our assay could in principle bind across two PD-1 peptides, the observed fast kinetics and low affinity were characteristic of single SH2 domain binding and not high affinity tandem SH2 binding, as for example observed for ZAP-70 and Syk in SPR (32, 33). Moreover, using PD-1 peptides with both ITIM and ITSM phosphorylated produced SPR traces similar to those with only the ITSM phosphorylated (data not shown). Therefore, SHP-1 and SHP-2 may differ in how they are recruited to the membrane (5, 34, 35).

### Molecular reach *in vivo*

Using the molecular reach for PD-1–SHP-1 complexes (14.6 nm) we showed that when uniformly distributed these complexes can only reach 15% of substrates. Given that PD-1 is a potent inhibitor of T cell activation, our results suggest other processes may be operating to increase its activity. One possible process is the induced co-clustering of PD-1 with substrates. We predict co-clustering can increase coverage to 90%, provided the density in these clusters is at least 10-fold higher than a uniform distribution. Indeed, microscopy experiments have found that inhibitory receptors that recruit SHP-1, including PD-1, form clusters with their substrates (5, 28, 36), although the precise density is presently unknown. Another possibility is that processes may operate to increase the molecular reach *in vivo*. In the case of other immune receptors, there is evidence that their tails can reversible associated with the plasma membrane and that this can be regulated (37–40), suggesting that the molecular reach may be dynamically regulated by signalling. This in turn suggests the possibility of molecular reach inhibitors that can control immune receptor activity by targeting the unstructured receptor tail or fleixble linkers within enzymes, which can have advantages over the targetting of structured domains (41–43)

### Tethering in signalling pathways

Although it is increasingly clear that cellular signalling relies on tethered reactions (3, 19, 44, 45), our quantitative understanding of this process remains limited. Mathematical models have shown how tethering increases intramolecular reactions rates by increasing local concentrations (20, 21). A feature of tethered reactions by immune receptors, and indeed many other membrane-confined reactions, is that they are intermolecular, meaning the substrate is a different receptor tail (i.e. a different tether). Although 2D intermolecular membrane reactions have been extensively studied, the models often simplify the reactions so that they take place within the 2D membrane rather then the 3D volume proximal to it (46–51). By explicitly modelling reach, it can be extracted from experimental data (as in the present work) and incorporated into accurate models of membrane reactions whereby membrane recruitment increases local concentrations but only if substrates are within reach. Such a model has recently demonstrated that increasing the molecular reach can both increase or decrease reactions rates depending on diffusion (24).

## Methods

### SHP-1 molecular reach estimates from structure

Using the structure of SHP-1 in the open conformation (PDB 3PS5) and sequence data from UniProt (P29350), we can estimate a range of reach values for SHP-1. Direct measurement from PDB structure of N-SH2 binding site to catalytic site gives a reach estimate of 5.3 nm. For our maximum reach estimate, we subdivide SHP-1 into 3 structured domains (N-SH2, C-SH2, PTP) and 2 linker domains. For the structured domains, distances were measured from the structure in PDB 3PS5: between binding pocket and linker for N-SH2, between two linkers for C-SH2, and from the linker to the active site for PTP. We then count the number of residues in the two intervening disordered linker domains, and compute contour length assuming these are fully extended. Adding these 5 numbers together (N-SH2, linker, C-SH2, linker, PTP) yields a value 20.4 nm. All measurements of structured domains were calculated using the measurement tool in PyMol.

### Peptides and SHP-1

All phosphopeptides were custom-synthesized by Peptide Protein Research, Ltd. and were N-terminally biotinylated. Peptide sequences are listed in Table 1. Human SHP-1 with an N-terminal 6x His tag was produced in *Escherichia coli* BL21-CodonPlus (DE3)-RIPL strain (Agilent Technologies) and purified on Ni^2+^-NTA agarose (Invitrogen) (16). Aliquots were stored at -80°C. On the day of experiment, SHP-1 was further purified by size-exclusion chromatography on an AKTA fast protein liquid chromatography system equipped with a Superdex S200 10/300 GL column (both from GE Healthcare Life Sciences) equilibrated with 10 mM Hepes (pH 7.4), 150 mM NaCl, 3 mM EDTA, 0.05% Tween 20 supplemented with 1 mM dithiothreitol. SHP-1 concentration was determined from absorbance at 280 nm measured on a Nanodrop ND-2000 spectrophotometer (Thermo Scientific).

**Table 1:** Peptides used in this study (phosphotyrosines are denoted as Y*)

| Name | Sequence |
| --- | --- |
| PEG28-PD-1 | biotin-(PEG)<sub>28</sub>-SVPEQTEY*ATIVFPSG |
| PEG12-PD-1 | biotin-(PEG)<sub>12</sub>-SVPEQTEY*ATIVFPSG |
| PEG6-PD-1 | biotin-(PEG)<sub>6</sub>-SVPEQTEY*ATIVFPSG |
| PEG3-PD-1 | biotin-(PEG)<sub>3</sub>-SVPEQTEY*ATIVFPSG |
| PEG0-PD-1 | biotin-SVPEQTEY*ATIVFPSG |
| PD-1 | biotin-SRAARGTIGARRTGQPLKEDPSAVPVFSVDYGELDFQ<br>WREKTPEPPVPSVPEQTEY*ATIVFPSG |
| SLAM | biotin-QLRRRGKTNHYQTTVEKKSLTIYAQVQKPGPLQKKLD<br>SFPAQDPCTTIYVAATEPVPESVQETNSITVY*ASVTLPES |

### Surface Plasmon Resonance

Experiments were performed on a Biacore T200 instrument (GE Healthcare Life Sciences) at 37°C and a flow rate of 10 *µ*l/min. Running buffer was the same as for size-exclusion chromatography. Streptavidin was coupled to a CM5 sensor chip using an amino coupling kit to near saturation, typically 10000–12000 response units (RU). Biotinylated peptides were injected into the experimental flow cells (FCs) for different lengths of time to produce desired immobilisation levels (typically 25–100 RU). Concentrations of immobilized peptides were determined from the RU values as described in (16). The molar ratio of peptide to streptavidin was kept below 0.25 to avoid generating streptavidin complexes with more than 1 peptide. Usually, FC1 and FC3 were used as references for FC2 and FC4, respectively. Excess streptavidin was blocked with biotin (Avidity). Before SHP-1 injection, the chip surface was conditioned with 10 injections of the running buffer and SHP-1 was then injected over all FCs; the duration of injections was the same for conditioning and SHP-1 injection (45 s).

### Solution assay for allosteric activation of SHP-1

The reaction mixture contained (final concentrations) 80 mM HEPES (pH 7.4), 1 mM DTT, 60 *µ*M PEG0-PD-1 peptide, 5% DMSO (vehicle), 10 mM p-nitrophenyl phosphate (*p*NPP), and 0.1 *µ*M SHP-1; the reaction was started by adding SHP-1. The reaction mixtures were incubated at 37°C. Aliquots were withdrawn at appropriate time points and dephosphorylation was stopped by addition of an equal volume of freshly prepared 80 mM HEPES (pH 7.4), 20 mM iodoacetamide, 100 mM Na_3_VO_4_. Absorbance at 405 nm was measured on the Nanodrop ND-2000. In the control, the quenching solution was added before SHP-1 and the mixture was kept either on ice or at 37°C for the duration of the time course. The efficiency of quenching was confirmed by the absence of a difference in absorbance between samples kept on ice or at 37°C.

### MPDPDE model and parameter fitting

We modified our previously reported multicenter particle density partial differential equation (MPDPDE) model (16). The original model included five fitting parameters (labelled *p*_1_ to *p*_5_) that are used to recover the five biophysical parameters: binding and unbinding rates *k*_on_, *k*_off_ of the enzyme to the receptor, the catalytic rate of the enzyme in solution *k*_cat_(solution), the catalytic rate of the enzyme when tethered to the receptor *k*_cat_(tethered), and the effective local concentration *σ*\* or equivalently, molecular reach of the reaction *L*. The present data exhibited nonspecific binding of the enzyme to the surface that differed in magnitude between the control and experimental flow cells and as a result, produced a different baseline before and after the SHP-1 injection. We modified the original model to include non-specific binding with rate *n*_ns_ that changed linearly with time (see Fig. S2) between the start (*p*_start_) and end (*p*_end_) of the SHP-1 injection. Therefore, the extended model included 3 additional fitted parameters compared to the original model (16).

To fit the SPR data to the extended MPDPDE model, we use a simulated annealing algorithm (52), with at least 10^5^ steps and a temperature function decreasing to zero as (1 – ([step]*/*10^5^)^4^). For initial guess, we used

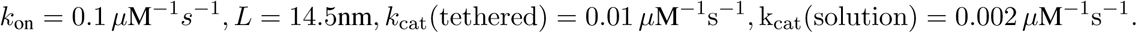

For the initial guess of *k*_off_, we first fit an exponential curve to the SPR time series data in the dissociation phase after SHP-1 injection ceases (e.g., after *t* = 45s in Fig. 2A). We find the parameters generated by simulated annealing are in close agreement with parameters found from MATLAB’s least-squares curve fitting (lsqcurvefit) function (not shown). However, the sum of square error for the parameters found using simulated annealing is consistently smaller. We perform simulated annealing three times on each dataset, using the fit with the lowest sum of square error for our analysis. To test for under-constrained parameter fitting, we perform a Markov Chain Monte Carlo (52) with unbounded, flat priors for each parameter. The resulting posteriors show that the parameters can be independently determined (Fig. S3). All model evaluation and fitting are implemented in MATLAB 2017b.

### Estimation of molecular reach

The molecular reach of the reaction, *L*, in our SPR assays is influenced by the reach for the tether, *L*_tether_, and the reach of the enzyme, *L*_SHP-1_. For a worm-like chain model, the probability density of a site on the molecule at location 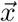 is

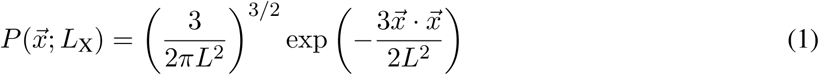

where *L*_X_ is a property of the molecule. For a worm-like chain model, 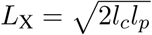, where *l*_*c*_ is the contour length and *l*_*p*_ is the persistence length, but we note that Eq. 1 arises in more general molecular models, so we use it to describe the behavior of the enzyme, without the interpretation of *L*_X_ in terms of a contour length and persistence length. In (16), we show that this leads to a local concentration kernel

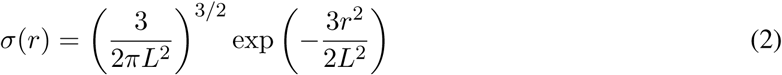

where *r* is the distance between the anchors of the two tethers, and

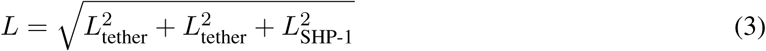

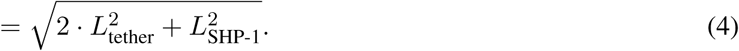

For disordered domains and PEG linkers, we interpret *L* in terms of the worm-like chain model (20, 53), so the reach can be estimated from the contour length and the persistence length of the domain. For the constructed PEG-PD-1 peptides, the contour length (from the surface anchor to binding site of SHP-1) is the number of PEG linkers *N*_PEG_ times the length of a single PEG, *l*_PEG_ ∼ 0.4 nm (54). From this, we derive an approximation for the reach of SHP-1,

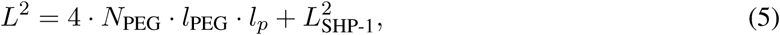

predicting that the reach is given by the intercept of the line *L*^2^ versus *N*_PEG_.

### Uncertainty quantification for derived parameters

For each PEG length, *L*^2^ is calculated by averaging the fitted parameter *σ*\* for all replicates and transforming the average to a single *L*^2^ value for the peptide. Error propagation is used to convert the standard deviation of *σ*\* to an error for *L*^2^.

Best fit lines with associated R-values and p-values for PEG28-PD1 parameters versus phosphatase and peptide concentrations, shown in Fig. 2 and 3 were determined using MATLAB’s robust fitlm function.

We use MATLAB’s anova1 and multcompare functions to conduct multiple comparison t-tests on paired PEG peptide parameters to establish significant differences. Pairs that are significantly different at the 0.05, 0.01, and 0.001 level are shown.

### Implications of reach for reactions at the cell membrane

If PD-1 is uniformly distributed on the cell, we estimate its surface density *ρ*_0_ by assuming 65,000 PD-1 molecules per T cell (55) and assuming that the T cell is approximately a sphere of radius 5 *µ*m, giving a PD-1 surface density of ∼2 · 10^−4^ nm^−2^. We use 1 *µ*M as the concentration of cytosolic SHP-1 (55). The effective concentration experienced by a substrate is then given by

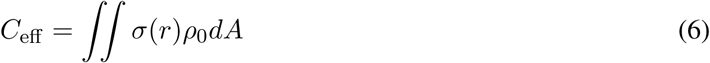

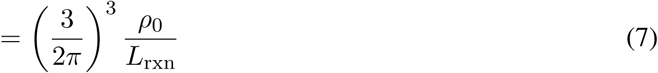

Since tethered reactions are heterogeneously distributed on the membrane, some substrate molecules are inaccessible to the enzyme. We determine what fraction of the substrate is accessible by a PD-1–SHP1 complex for a given reach of reaction and PD-1 density. To do this, we calculate the probability of at least one PD-1–SHP1 complex being within a circle of radius L, of any given substrate. We assume finding a number of activated, SHP-1-bound PD-1 molecules within a disk around the substrate is Poisson distributed, with 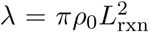 and *ρ*_0_ estimated above. We use this to determine the probability of at least one PD-1– SHP1 complex within reach,

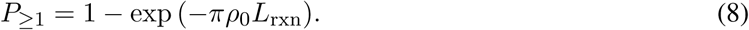

## Acknowledgements

We thank Marion H. Brown, P. Anton van der Merwe, and members of the van der Merwe and Dushek laboratories for helpful discussions. The work has been funded by NSF CAREER grant (DMS 1454739 to JA), NSF grant DMS 1763272 for funding JA and LC and a grant from the Simons Foundation (594598, QN) for funding JA and LC, and a Wellcome Trust Senior Research Fellowship (207537/Z/17/Z to OD).

## Supplementary Information

**Table S1:**
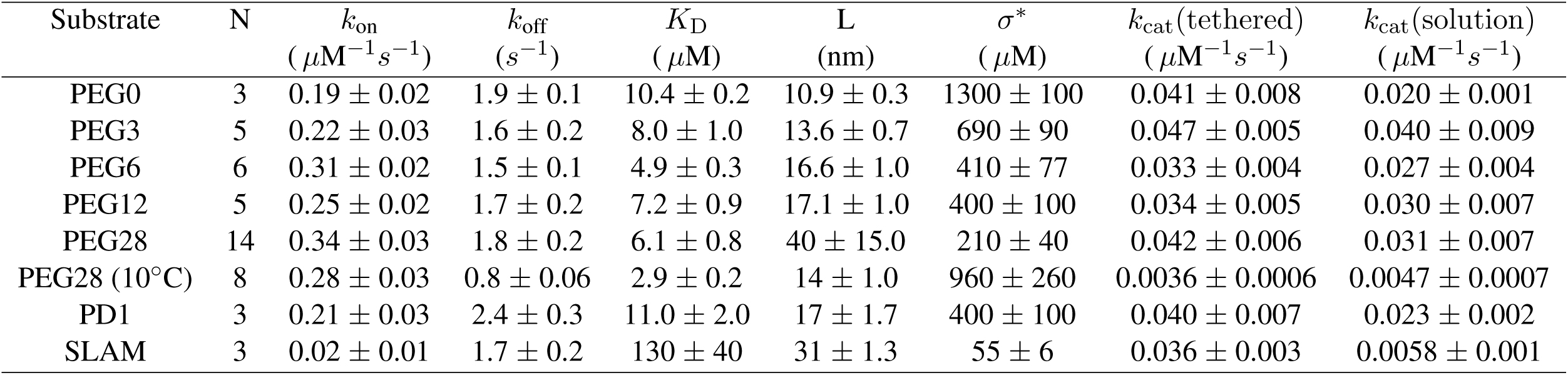
Average biophysical parameter values for each peptide. All experiments conducted at temperature 37°C except where noted.

| Substrate | N | $k_{\text{on}}$<br>($\mu\text{M}^{-1}\text{s}^{-1}$) | $k_{\text{off}}$<br>($\text{s}^{-1}$) | $K_D$<br>($\mu\text{M}$) | L<br>(nm) | $\sigma^*$<br>($\mu\text{M}$) | $k_{\text{cat}}$(tethered)<br>($\mu\text{M}^{-1}\text{s}^{-1}$) | $k_{\text{cat}}$(solution)<br>($\mu\text{M}^{-1}\text{s}^{-1}$) |
| --- | --- | --- | --- | --- | --- | --- | --- | --- |
| PEG0 | 3 | $0.19 \pm 0.02$ | $1.9 \pm 0.1$ | $10.4 \pm 0.2$ | $10.9 \pm 0.3$ | $1300 \pm 100$ | $0.041 \pm 0.008$ | $0.020 \pm 0.001$ |
| PEG3 | 5 | $0.22 \pm 0.03$ | $1.6 \pm 0.2$ | $8.0 \pm 1.0$ | $13.6 \pm 0.7$ | $690 \pm 90$ | $0.047 \pm 0.005$ | $0.040 \pm 0.009$ |
| PEG6 | 6 | $0.31 \pm 0.02$ | $1.5 \pm 0.1$ | $4.9 \pm 0.3$ | $16.6 \pm 1.0$ | $410 \pm 77$ | $0.033 \pm 0.004$ | $0.027 \pm 0.004$ |
| PEG12 | 5 | $0.25 \pm 0.02$ | $1.7 \pm 0.2$ | $7.2 \pm 0.9$ | $17.1 \pm 1.0$ | $400 \pm 100$ | $0.034 \pm 0.005$ | $0.030 \pm 0.007$ |
| PEG28 | 14 | $0.34 \pm 0.03$ | $1.8 \pm 0.2$ | $6.1 \pm 0.8$ | $40 \pm 15.0$ | $210 \pm 40$ | $0.042 \pm 0.006$ | $0.031 \pm 0.007$ |
| PEG28 (10°C) | 8 | $0.28 \pm 0.03$ | $0.8 \pm 0.06$ | $2.9 \pm 0.2$ | $14 \pm 1.0$ | $960 \pm 260$ | $0.0036 \pm 0.0006$ | $0.0047 \pm 0.0007$ |
| PD1 | 3 | $0.21 \pm 0.03$ | $2.4 \pm 0.3$ | $11.0 \pm 2.0$ | $17 \pm 1.7$ | $400 \pm 100$ | $0.040 \pm 0.007$ | $0.023 \pm 0.002$ |
| SLAM | 3 | $0.02 \pm 0.01$ | $1.7 \pm 0.2$ | $130 \pm 40$ | $31 \pm 1.3$ | $55 \pm 6$ | $0.036 \pm 0.003$ | $0.0058 \pm 0.001$ |

**Figure S1:**
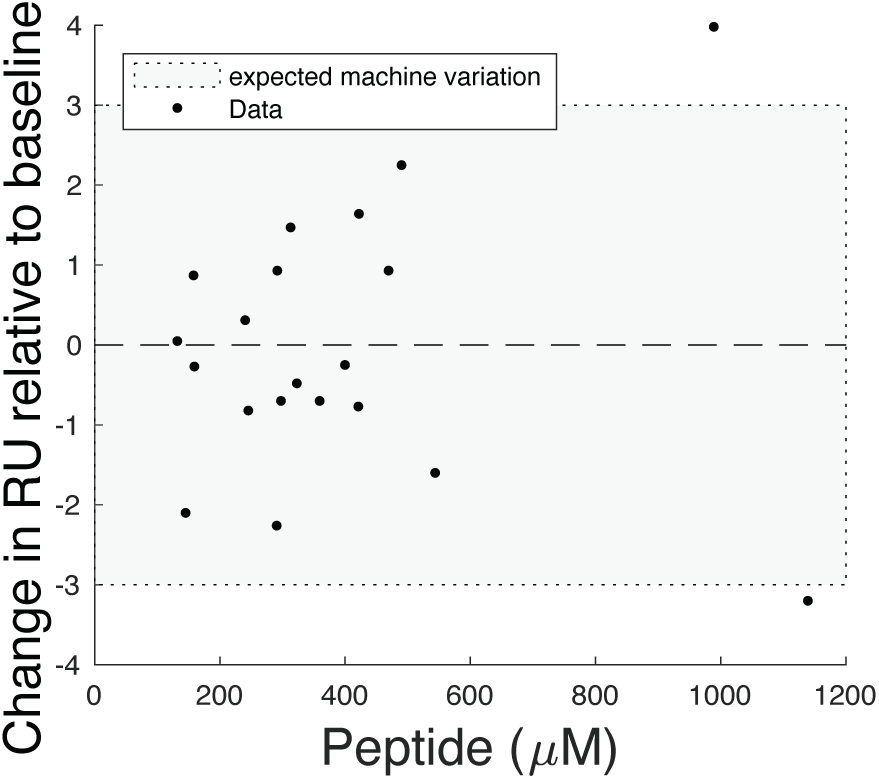
Change in baseline before and after phosphatase injection is not correlated with phosphorylated peptide concentration. The observed reduction in baseline RU after SHP-1 injection (see Fig. 2A,B) could be a result of the loss of phosphate mass from the chip surface. To investigate this possibility, alkaline phosphatase was injected over a surface immobilised with the indicated concentration of phosphorylated PEG28-PD-1 peptide for 300 seconds and the difference in baseline RU before and after injection was calculated. However, no correlation was observed with peptide concentration. The small ∼3 RU deviation is consistent with machine drift which is reported to be in the range of ∼1 RU per 100 seconds (gray area for our 300 second injection).

**Figure S2:**
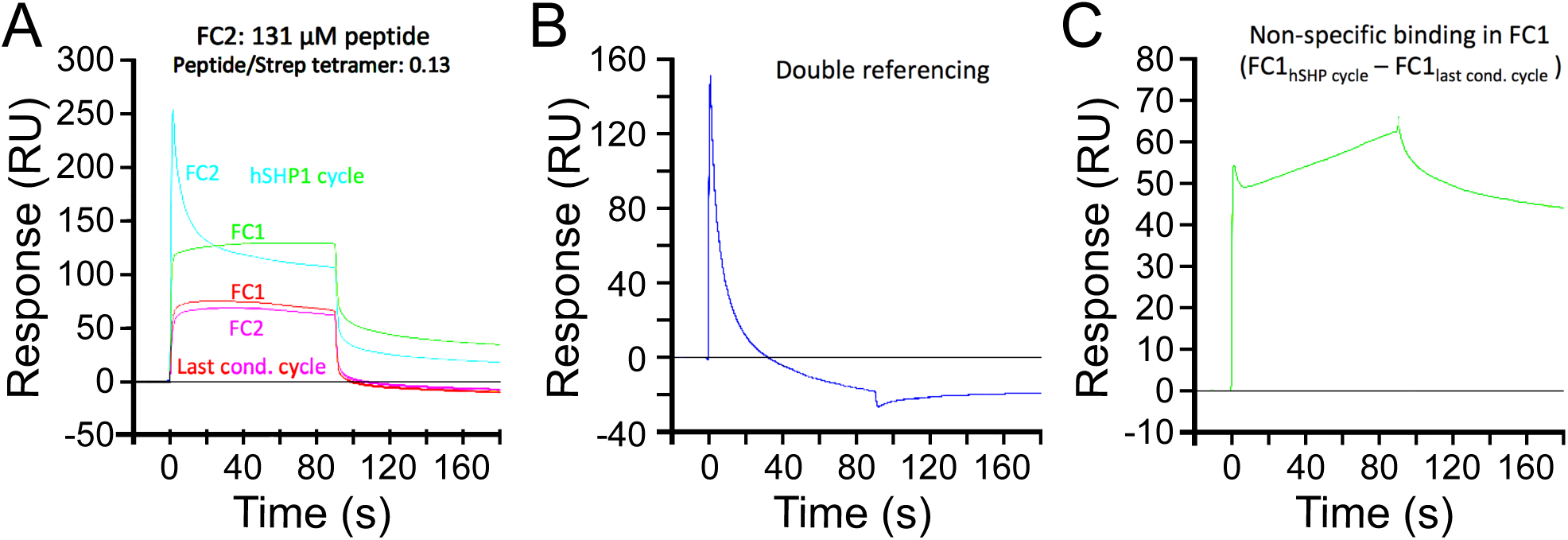
Change in baseline before and after SHP-1 injection can be explained by non-specific SHP-1 binding. (A) SPR traces of buffer injection (conditioning cycle) over flow cells 1 (red) and 2 (pink) followed by SHP-1 injection over flow cell 1 (green) and 2 (cyan). Flow cell 1 is a blank control whereas PEG28-PD-1 is immobilised in flow cell 2. All experiments were at at 37°C. Non-specific binding is apparent after SHP-1 injection because baseline remains above zero. (B) Double-referenced SPR trace for PEG28-PD-1 highlights a negative baseline after SHP-1 injection. This result suggests that non-specific binding in the control and experimental flow cells is not exactly matched. (C) Subtraction of the buffer injection from the SHP-1 injection in the control flow cell demonstrates that non-specific binding of SHP-1 can be approximated to be linear in time.

**Figure S3:**
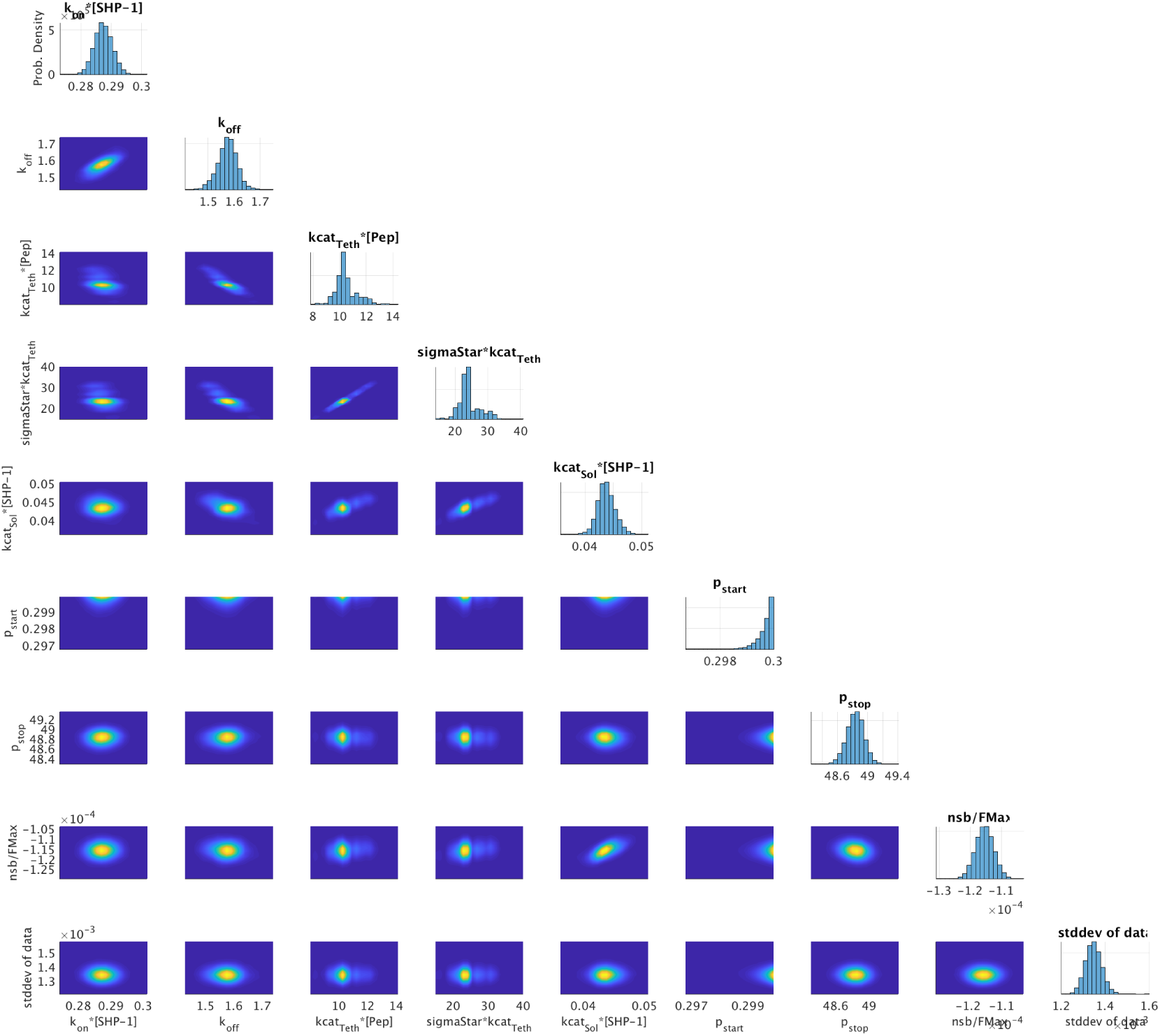
Posterior distributions of individual fitted parameters. Using Markov Chain Monte Carlo to fit the MPDPDE model to data obtained at 1 *µ*M SHP-1 and 125 *µ*M PEG28, demonstrates parameter estimates are well-identified.

**Figure S4:**
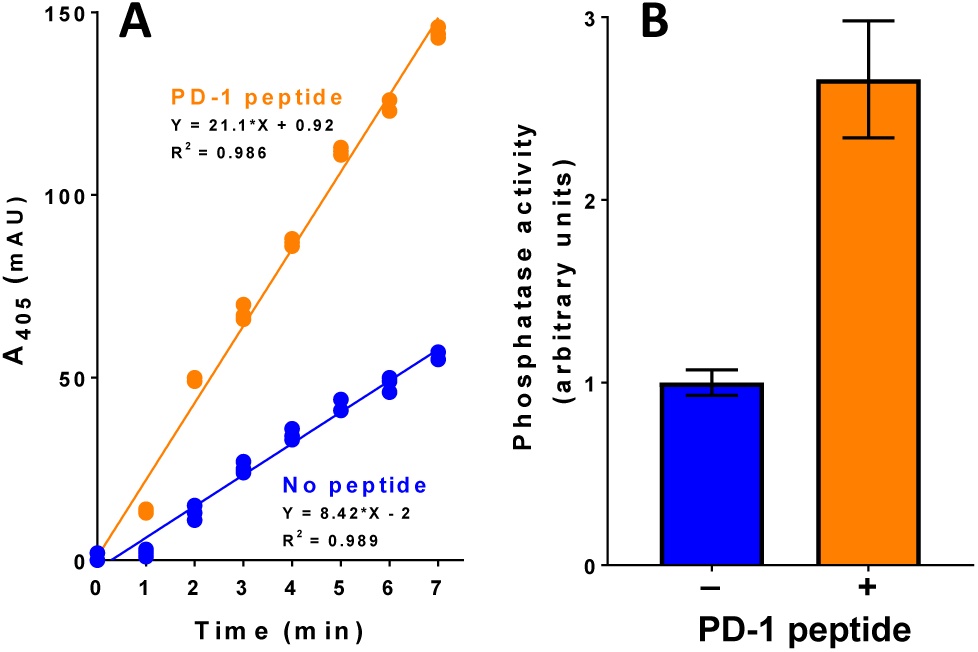
Solution-based assay for SHP-1 catalytic activity confirms a modest allosteric activation by singly phosphorylated PD-1 peptide. Human SHP-1 (0.1 *µ*M) was incubated at 37°C with 10 mM pNPP in the absence (blue) or presence (orange) of 60 *µ*M PEG0-PD-1 peptide. (A) Time course of pNPP dephosphorylation measured by absorbance at 405 nm at the indicated time points. Data points are technical replicates (n = 3). (B) The catalytic activity is determined by the fold-change in the slope of the time course data with a mean of 2.7. Error bars represent minimum and maximum from independent experiments (n=2).

**Figure S5:**
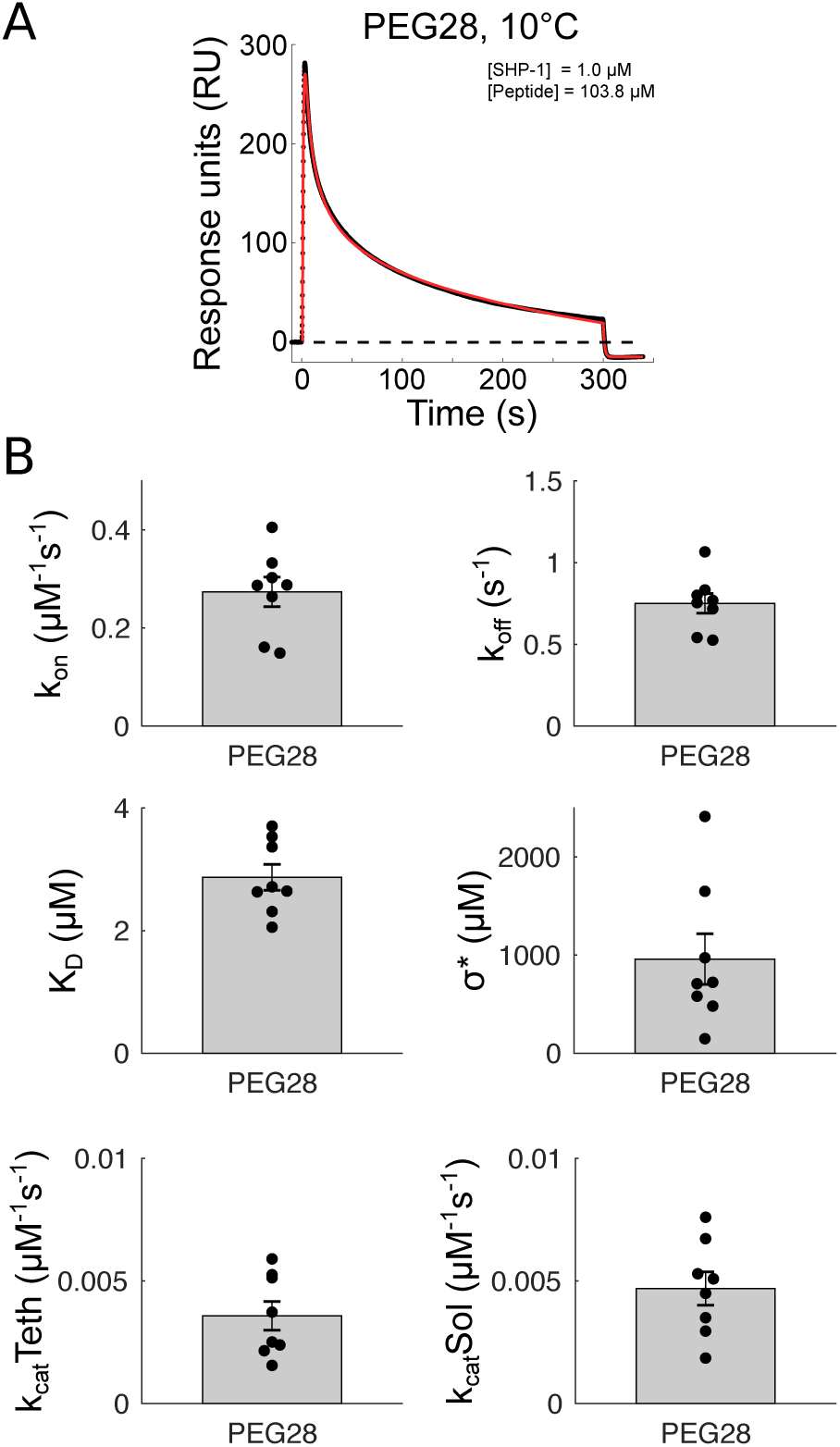
SHP-1 injection over PEG28-PD-1 at 10°C. (A) Representative SPR trace (black dots) and extended MPDPDE model fit (red line). (B) Averages and SEMs (gray with black error bars). Individual data points plotted as black dots.

**Figure S6:**
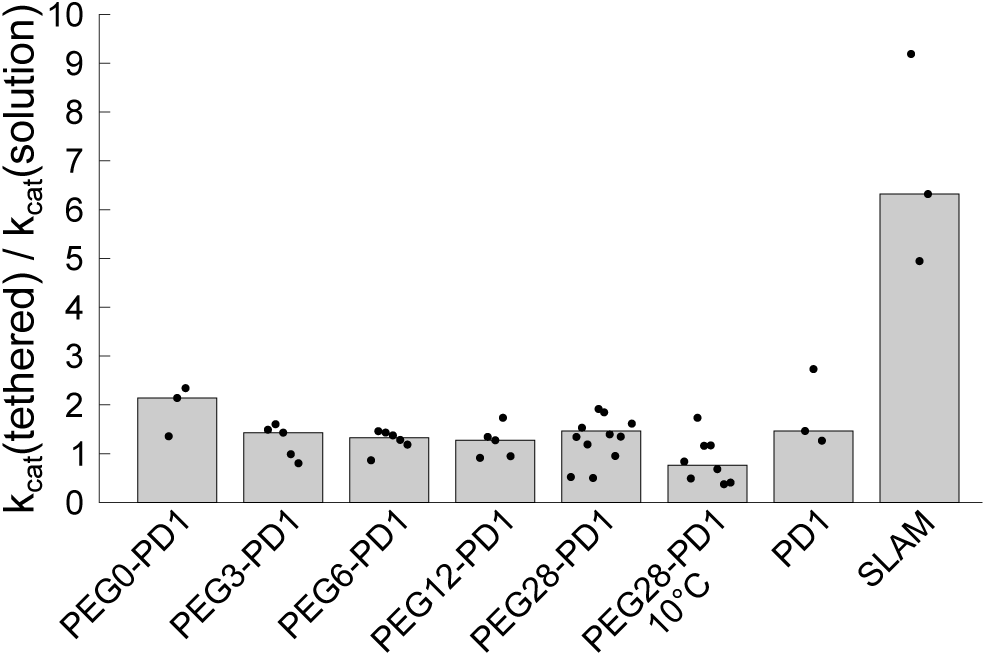
Summary of *k*_cat_(tethered)/*k*_cat_(solution) for all experimental conditions. Ratio of tethered to solution catalysis (*k*_cat_(tethered)/*k*_cat_(solution)) determined by SPR for the indicated conditions (all experiments at 37°C except where indicated). The median (gray bars) reveals only modest ∼1-2-fold allosteric activation for PD-1 across all conditions but a larger 6.2-fold activation for SLAM, which is discussed in the main text.

